# XerC is required for the repair of antibiotic- and immune-mediated DNA damage in *Staphylococcus aureus*

**DOI:** 10.1101/2022.09.04.506505

**Authors:** Elizabeth V. K. Ledger, Katie Lau, Edward W. Tate, Andrew M. Edwards

## Abstract

To survive in the host environment, pathogenic bacteria need to be able repair DNA damage caused by both antibiotics and the immune system. The SOS response is a key bacterial pathway to repair DNA double strand breaks and may therefore be a good target for novel therapeutics to sensitise bacteria to antibiotics and the immune response. However, the genes required for the SOS response in *Staphylococcus aureus* have not been fully established. Therefore, we carried out a screen of mutants involved in various DNA repair pathways to understand which were required for induction of the SOS response. This led to the identification of 16 genes that may play a role in SOS response induction, and of these, 3 that affected susceptibility of *S. aureus* to ciprofloxacin. Further characterisation revealed that, in addition to ciprofloxacin, loss of the tyrosine recombinase XerC increased the susceptibility of *S. aureus* to various classes of antibiotics, as well as to host immune defences. Therefore, the inhibition of XerC may be a viable therapeutic approach to sensitise *S. aureus* to both antibiotics and the immune response.

## Introduction

*Staphylococcus aureus* is a leading cause of infections, ranging from those affecting the skin and soft tissues to life-threatening invasive diseases such as bacteraemia and endocarditis [Turner *et al*., 2019]. Treatment of these infections can be challenging, especially when they are caused by multi-drug resistant strains. However, even in the case of infections caused by drug-susceptible strains, high rates of relapse and development of chronic infections are observed [Chang *et al*., 2003; Choi *et al*., 2021].

For *S. aureus* to survive in the host and establish an infection it must be able to survive damage caused by both the immune system and antibiotics. One of the main immune cell types used by the host to control *S. aureus* infections is neutrophils, which typically kill *S. aureus* via antimicrobial peptides and reactive oxygen species (ROS) released during the oxidative burst [Rigby *et al*., 2012]. These ROS damage many cellular components, including nucleic acids, which results in DNA double strand breaks that are lethal if not repaired by the bacterium [Nguyen *et al*., 2017; Li *et al*., 2021]. Some clinically important antibiotics directly target DNA synthesis/ replication, for example fluoroquinolones, which inhibit DNA gyrase, or co-trimoxazole, which inhibits tetrahydrofolate synthesis, thereby causing stalled DNA replication. In addition, other classes of bactericidal antibiotic, including beta lactams and daptomycin, cause DNA damage indirectly via the production of ROS [Kohanski *et al*., 2007; Po *et al*., 2021; Clarke *et al*., 2021]. Therefore, it is well established that *S. aureus* needs to be able to repair DNA damage to survive inside the host.

The main pathway used by bacteria to repair DNA damage is the SOS response, which regulates expression of genes responsible for DNA repair [Ha and Edwards, 2021]. In *S. aureus*, DNA double strand breaks are processed by the helicase/nuclease activity of RexAB (a member of the RecBCD/AddAB family) to generate single stranded DNA [Ha *et al*., 2020]. This is then bound by RecA, forming a nucleoprotein filament that triggers the autocleavage of LexA, a transcriptional repressor, derepressing expression of genes in the SOS regulon [Ha and Edwards, 2021]. In *S. aureus*, the SOS regulon consists of 16 genes, including *lexA* and *recA*, and several genes involved in repairing various types of DNA damage, including nucleotide excision repair and the processing of stalled replication forks [Cirz *et al*., 2007]. Many antibiotics have been shown to induce the SOS response in *S. aureus*, including directly DNA damaging agents such as ciprofloxacin, co-trimoxazole and nitrofurantoin but also antibiotics with other targets, including daptomycin, oxacillin and chloramphenicol [Clarke *et al*., 2019; Cirz *et al*., 2007; Maiques *et al*., 2006; Clarke *et al*., 2021]. In line with this, a mutant defective for *rexB* was unable to induce the SOS response when exposed to these agents and showed increased antibiotic susceptibility [Clarke *et al*., 2021].

In addition to regulating DNA repair, the SOS response plays a key role in the emergence of antibiotic resistance. One of the genes most strongly expressed in response to induction of the SOS response is *umuC*, which encodes an error prone polymerase [Cirz *et al*., 2007; Sale *et al*., 2012]. Therefore, activation of the SOS response leads to an increase in the mutation rate, promoting the emergence of antibiotic resistance. As well as this, induction of the SOS response activates prophages present in the *S. aureus* genome, leading to the dissemination of virulence and antibiotic resistance genes via horizontal gene transfer [Ubeda *et al*., 2005].

Since the SOS response promotes survival of both antibiotics and the immune response as well as increasing the emergence of antibiotic resistance and the spread of virulence genes, its inhibition is an attractive therapeutic target to sensitise *S. aureus* to the action of antibiotics and the immune response. In support of this, several inhibitors of RecBCD/AddAB family proteins have been identified, including the Gam protein of bacteriophage lambda which sensitises *E. coli* to ciprofloxacin [Wilkinson *et al*., 2016; Lanyon-Hogg *et al*., 2021]. However, due to poor pharmacokinetic/pharmacodynamic properties and/or toxicity none are suitable therapeutics.

Unfortunately, while the SOS response has been well characterised in several model bacteria, the response of *S. aureus* differs significantly from these and is poorly understood. For example, the SOS regulons of *E. coli* and *B. subtilis* comprise 43 and 63, genes, respectively, many more than the 16 in *S. aureus* [Ha and Edwards, 2021]. Additionally, which genes are required for induction of the SOS response, and may therefore constitute good therapeutic targets, is not well understood. Therefore, to address this, we carried out a screen of genes previously identified to play a role in DNA repair, to determine which were important for inducing the SOS response and modulating susceptibility to antibiotics and the immune response.

## Results

### XerC and XseA are required for the repair of ciprofloxacin-mediated DNA damage

To establish which *S. aureus* genes were required for the SOS response, we identified and screened 54 mutants from the NARSA transposon library with transposons inserted in genes that had previously been described to play or proposed to play a role in DNA repair (Table S1) [Fey *et al*., 2013]. Where two genes contributed to a single protein complex (e.g. RexAB), only one gene was included in the screen. To measure the SOS response in each of these mutants, a previously well-characterised reporter plasmid consisting of the *recA* promoter upstream of *gfp* was transduced into the JE2 WT strain and each transposon mutant [Clarke *et al*., 2019]. Strains were then exposed to a range of therapeutically-relevant concentrations of the DNA-damaging antibiotic, ciprofloxacin (0 – 16 μg ml^-1^) and the SOS response measured over time (Fig. S1). The peak fluorescence value of each strain at 16 μg ml^-1^ ciprofloxacin was plotted to enable comparison between strains (Fig. 1A). This led to the identification of 16 mutants with a significantly reduced SOS response compared to the WT strain, including the *rexB*::Tn mutant, which has been previously identified as essential for induction of the SOS response in *S. aureus* and thus validated our approach.

**Figure 1.**
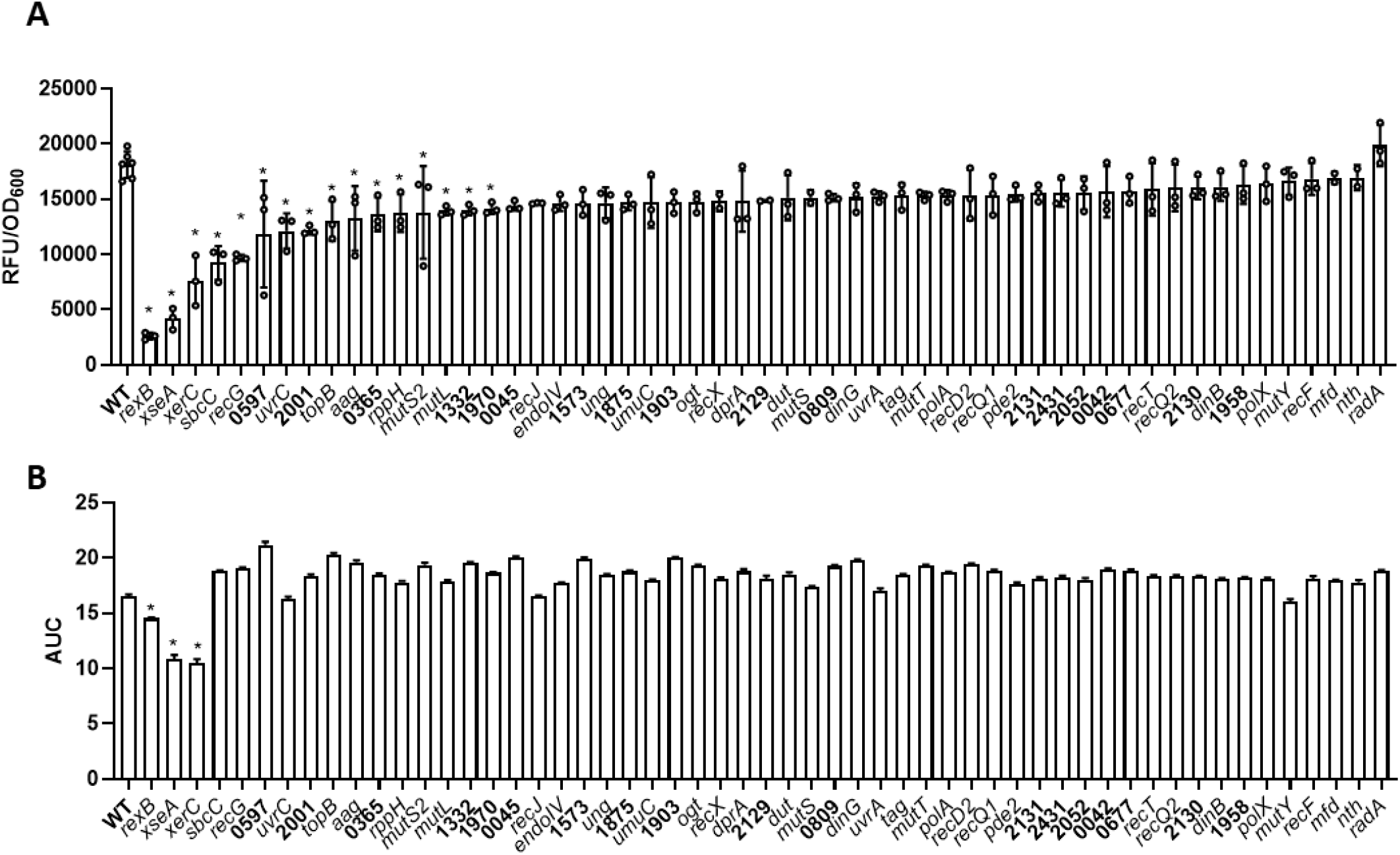
XerC and XseA are required for repair of ciprofloxacin-mediated DNA damage. *S. aureus* USA300 JE2 WT and mutants defective in various DNA repair genes containing the P*recA-gfp* SOS response reporter plasmid were exposed to 16 μg ml^-1^ ciprofloxacin for 17 h. (**A**) Induction of the SOS response was determined through measuring GFP fluorescence and the peak fluorescence value of each strain (adjusted for cell density) was plotted. (**B**) OD_600_ was measured every 15 min throughout the 17 h ciprofloxacin exposure and area under the curve plotted. Data in **A** represent the mean ± standard deviation of three independent experiments and data in **B** represent the mean ± standard error of three independent experiments. In each case, data were analysed by one-way ANOVA with Dunnett’s *post-hoc* test (* P < 0.05; WT vs mutant).

Secondly, we generated growth curves of each strain during a 17 h exposure to 16 μg ml^-1^ ciprofloxacin and plotted the area under the curve to give a measure of the susceptibility of each mutant to the antibiotic (Fig. S2). This demonstrated that three strains, *rexB*::Tn, *xerC*::Tn and *xseA*::Tn, were more susceptible to ciprofloxacin than the WT strain (Fig. 1B).

Taken together, we identified 16 mutants with an impaired ability to induce the SOS response on exposure to ciprofloxacin, and of these, three mutants were also more susceptible to ciprofloxacin, suggesting that they may be good targets to sensitise *S. aureus* to DNA-damaging antibiotics.

### XerC is required for the SOS response induced by various DNA-damaging agents

In addition to ciprofloxacin, many other DNA-damaging agents also induce the SOS response. Therefore, we next aimed to determine whether the impaired SOS induction occurred in response to various agents which cause DNA damage via diverse mechanisms or whether it was unique to ciprofloxacin. As we have previously demonstrated that RexB is required for induction of the SOS response, and preliminary experiments with the *xseA*::Tn strain produced inconsistent results, we focused on the role of XerC [Ha *et al*., 2020].

To do this, we exposed the WT and *xerC*::Tn mutant to a range of concentrations of ciprofloxacin, co-trimoxazole, nitrofurantoin, mitomycin C or paraquat, which generates superoxide radicals, whilst measuring the SOS response using the P*recA-gfp* reporter system. In each case, a dose-dependent increase in induction of the SOS response was observed in the WT strain (Fig. 2A – E). However, with the exception of nitrofurantoin, the SOS response of the *xerC*::Tn mutant strain was significantly lower than that of the WT strain at higher concentrations of the genotoxic agents, indicating that XerC is required for maximal SOS induction in response to diverse types of DNA-damage (Fig. 2A – E).

**Figure 2.**
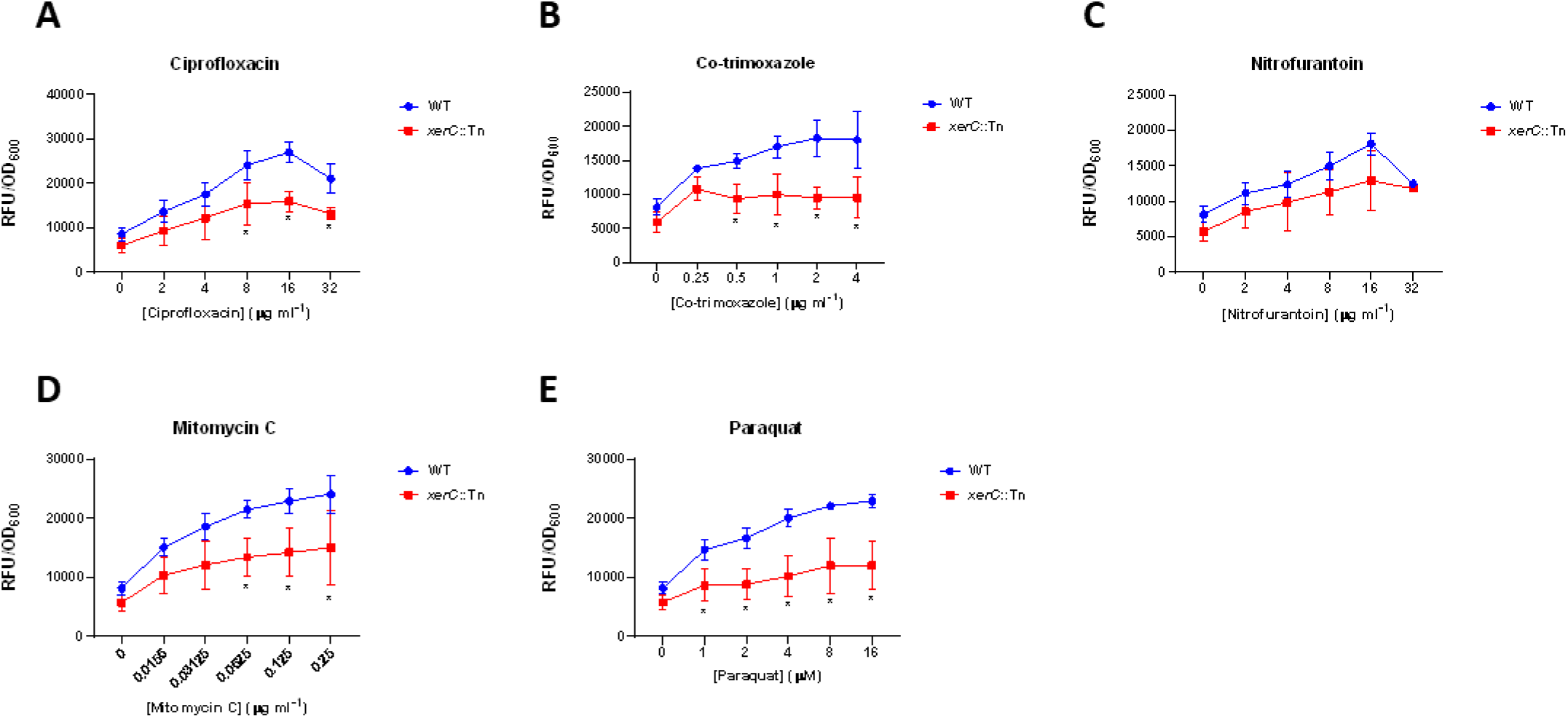
XerC is required for SOS response induced by various DNA-damaging agents. *S. aureus* USA300 JE2 WT and the *xerC*::Tn mutant containing the P*recA-gfp* SOS response reporter plasmid were exposed to indicated concentrations of (A) ciprofloxacin, (B) co-trimoxazole, (C) nitrofurantoin, (D) mitomycin C or (E) paraquat before the peak fluorescence value of each strain (adjusted for cell density) was plotted. Data represent the mean ± standard deviation of three independent experiments and were analysed by two-way ANOVA with Dunnett’s *post-hoc* test (* P < 0.05; WT vs mutant at indicated concentration).

In many species of bacteria, XerC has been implicated in the monomerisation of plasmids, enabling the stable segregation of plasmids during cell division [Sirois *et al*., 1995]. As our reporter system was plasmid based, we checked whether an alternative explanation for the reduced SOS response of the mutant strain was a loss of the reporter system during the assay. To do this, we carried out a passage experiment, repeatedly subculturing both the WT and *xerC*::Tn strains carrying the P*recA-gfp* reporter plasmid in either the presence of kanamycin, to select for the plasmid, or the absence of selection pressure, to replicate the conditions of the SOS reporter assay. Each passage consisted of a 1:1000 dilution of the bacterial cultures, enabling approximately 10 cell divisions each passage. This was repeated 6 times, resulting in approximately 60 generations occurring during the assay. As expected, in the presence of kanamycin, there was no significant loss of plasmid throughout the experiments in either the WT or the *xerC*::Tn strain (Fig. 3). By contrast, in the absence of selection, the plasmid was slowly lost from the WT strain, while between 20 and 30 generations, there was a significant reduction of the number of *xerC*::Tn bacteria that carried the plasmid, with over 95 % of colonies having lost the plasmid by 30 generations (Fig. 3). However, despite this, this plasmid loss would not be expected to explain the loss of SOS response, as this assay was completed within four generations.

**Figure 3.**
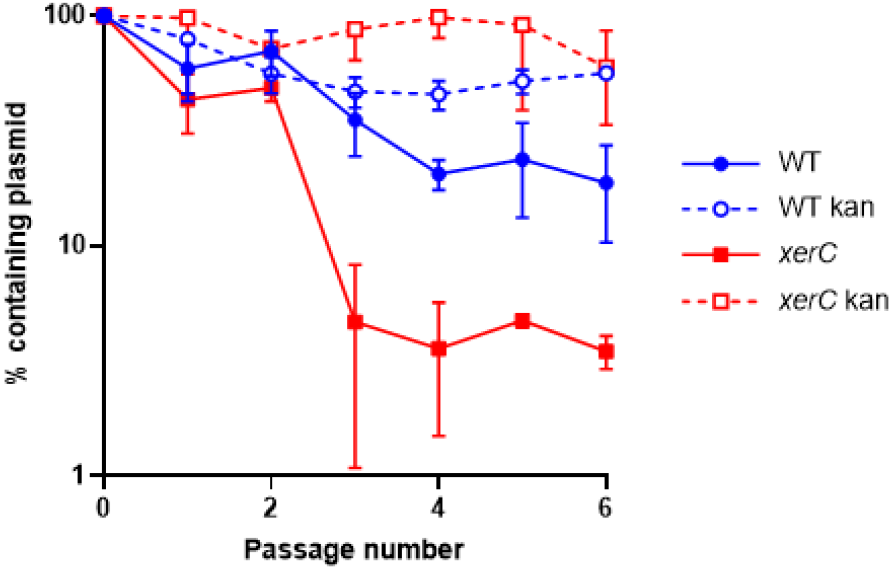
Loss of reporter plasmid does not explain reduced SOS response. *S. aureus* USA300 JE2 WT and *xerC*::Tn containing the P*recA-gfp* reporter plasmids underwent 6 passages in TSB or TSB supplemented with 90 μg ml^-1^ kanamycin. After each passage, strains were plated on TSA and TSA supplemented with 90 μg ml^-1^ kanamycin and the percentage of colonies containing the plasmid determined. Data represent the mean ± standard deviation of three independent experiments.

### Lack of XerC-mediated DNA repair sensitises *S. aureus* to various classes of antibiotic

Next, we aimed to determine whether XerC-mediated DNA repair affected susceptibility of *S. aureus* to a range of antibiotics. We measured the susceptibility of the WT and *xerC*::Tn strains to a panel of antibiotics, including several which directly damage DNA, as well as cell wall targeting antibiotics and protein synthesis inhibitors, some of which have been reported to damage DNA indirectly, for example via the production of ROS.

Susceptibility to four directly DNA-damaging antimicrobials was determined by establishing their minimum inhibitory concentrations, ciprofloxacin, co-trimoxazole, nitrofurantoin and mitomycin C. In each case, the *xerC*::Tn mutant was more susceptible than WT, with the largest increases in susceptibility (8-fold) being observed with co-trimoxazole and mitomycin C (Fig. 4). In addition, the *xerC*::Tn mutant was also more susceptible to the cell wall synthesis inhibitors fosfomycin and cloxacillin, but not daptomycin or vancomycin, which target the cell envelope, and each of the four protein synthesis inhibitors examined, gentamicin, tetracycline, rifampicin and chloramphenicol (Fig. 4).

**Figure 4.**
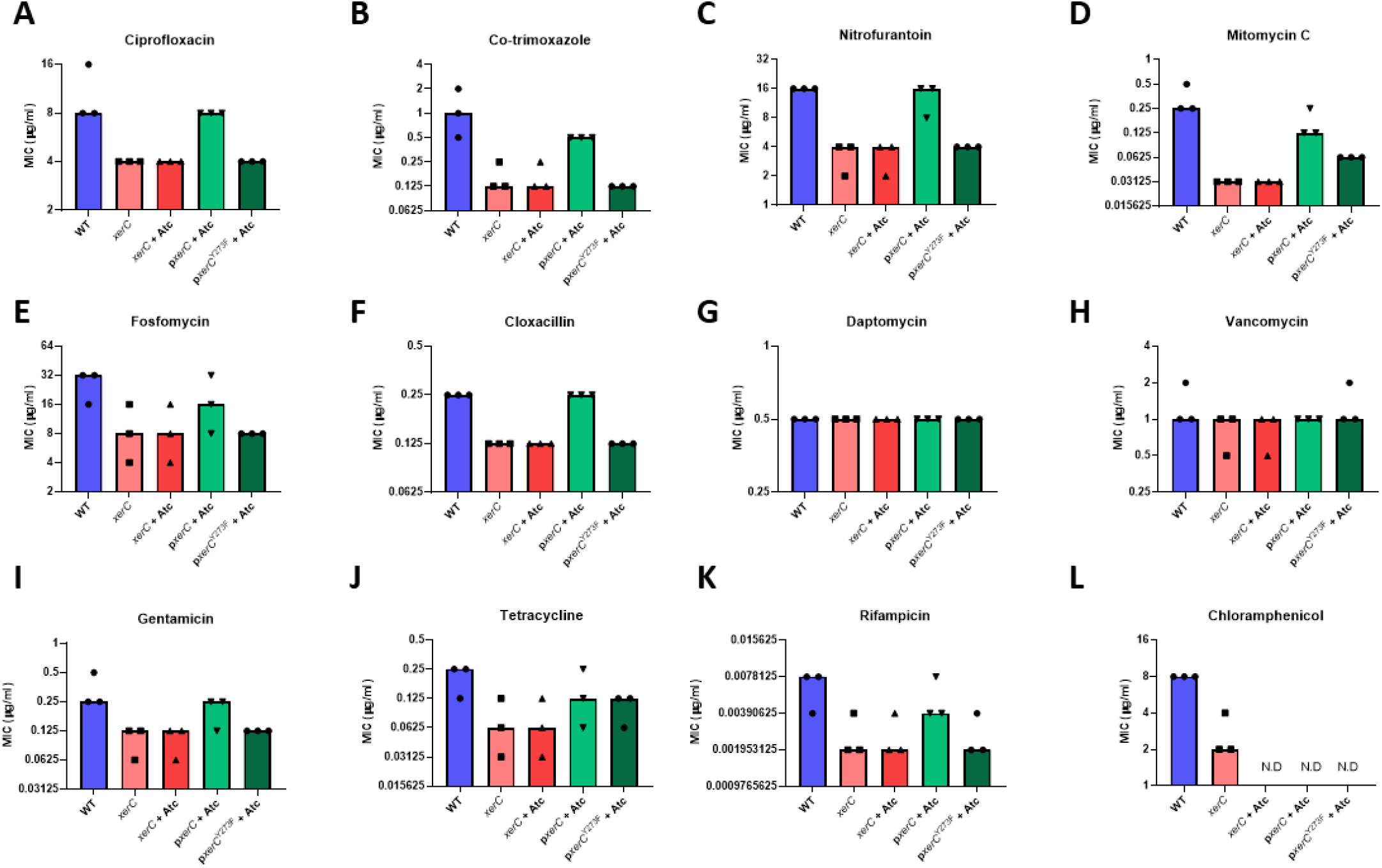
Lack of XerC-mediated DNA repair sensitises *S. aureus* to various classes of antibiotic. MICs of *S. aureus* USA300 JE2 WT, the *xerC*::Tn mutant and the *xerC*::Tn mutant complemented with p*xerC* or p*xerC*^Y273F^ to (**A**) ciprofloxacin, (**B**) co-trimoxazole, (**C**) nitrofurantoin, (**D**) mitomycin C, (**E**) fosfomycin, (**F**) cloxacillin, (**G**) daptomycin, (**H**) vancomycin, (**I**) gentamicin, (**J**) tetracycline, (**K**) rifampicin and (**L**) chloramphenicol. Data represent the median of three independent experiments. N.D, not determined; Atc, anhydrotetracycline.

Next, we aimed to confirm that this was due to the DNA-repair ability of XerC. XerC is a tyrosine-type site-specific recombinase, which uses an active site tyrosine (Y273) to break and rejoin DNA at a specific sequence [Hallet *et al*., 1999]. To do this, we complemented *xerC*::Tn with either WT *xerC* or *xerC*^Y273F^, a catalytically inactive version with a mutated active site tyrosine to abolish the recombination activity of XerC, to under the control of a tetracycline-inducible promoter [Gründling and Schneewind, 2007]. The plasmid used for complementation conferred chloramphenicol resistance, precluding further testing of whether XerC contributed to chloramphenicol susceptibility. In each case, complementation with the WT copy of *xerC* at least partially restored antibiotic susceptibility to WT levels (Fig. 4). However, complementation with *xerC*^Y273F^ did not, indicating that the increased sensitivity of the *xerC*::Tn mutant was due to its lack of DNA repair ability (Fig. 4).

Finally, we measured the survival of bacteria exposed to antibiotics as a second measure of drug susceptibility. To do this, 10^8^ CFU ml^-1^ of each strain were exposed to 5 x the MIC of the WT strain for 6 h before surviving bacteria were enumerated. In agreement with the MIC data, the *xerC*::Tn mutant was killed to a greater extent than the WT strain on exposure to each of the directly DNA-damaging antibiotics (Fig. 5). Similarly, greater killing of the mutant strain was observed on exposure to fosfomycin and cloxacillin as well (Fig. 5). In line with the observation that the WT and mutant strains had the same MICs to vancomycin and daptomycin, no differences in killing at 6 h were seen with these antibiotics (Fig. 5). However, in contrast to the MIC data, no differences in survival between the strains were observed with any of the protein synthesis inhibitors (Fig. 5).

**Figure 5.**
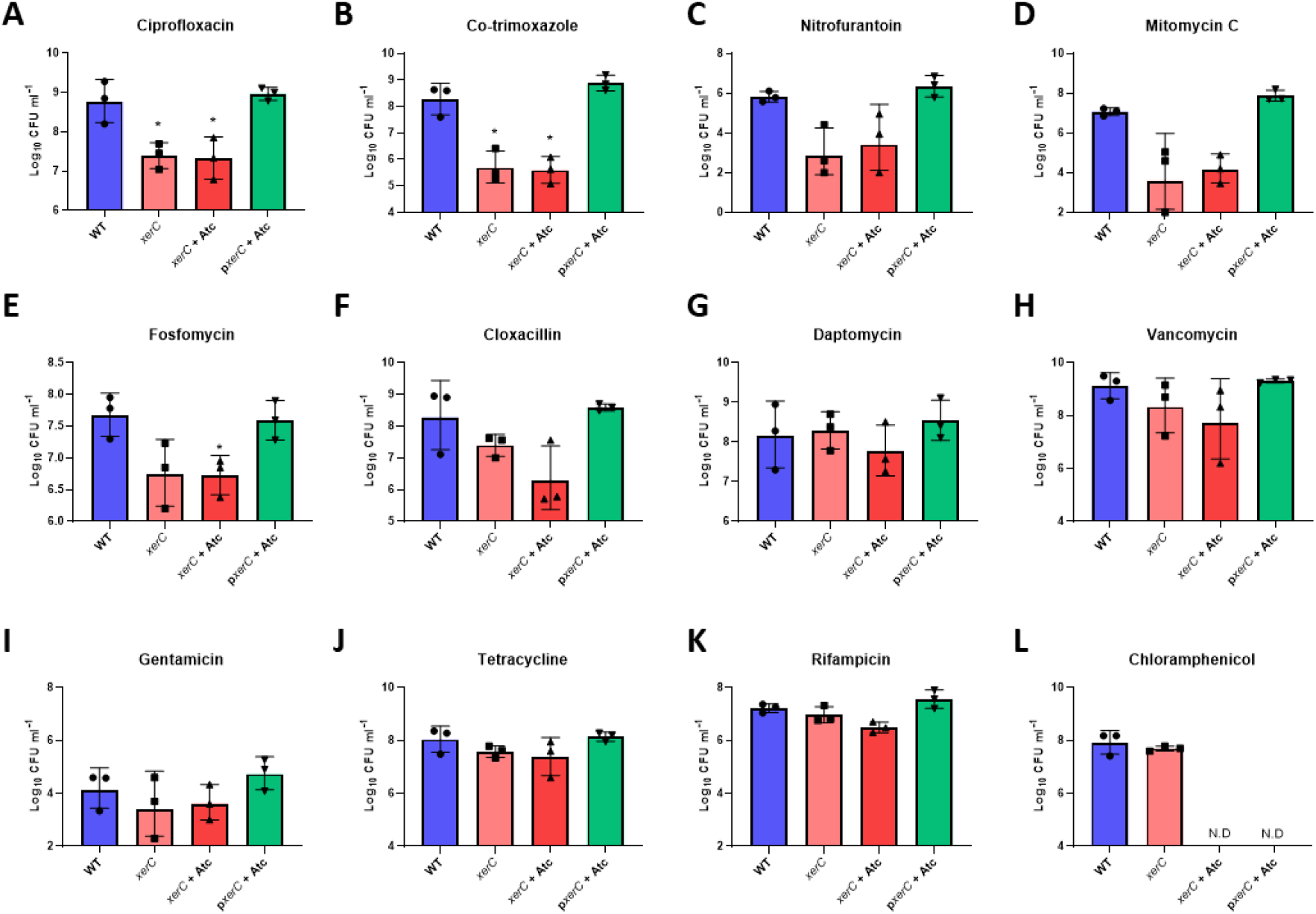
Lack of XerC sensitises *S. aureus* to various classes of antibiotic. Log_10_ CFU ml^-1^ of *S. aureus* USA300 JE2 WT, the *xerC*::Tn mutant and the *xerC*::Tn mutant complemented with p*xerC* or p*xerC*^Y271F^ after a 6 h exposure of 10^8^ CFU ml^-1^ to (**A**) 40 μg ml^-1^ ciprofloxacin, (**B**) 5 μg ml^-1^ co-trimoxazole, (**C**) 80 μg ml^-1^ nitrofurantoin, (**D**) 1.25 μg ml^-1^ mitomycin C, (**E**) 160 μg ml^-1^ fosfomycin, (**F**) 1.25 μg ml^-1^ cloxacillin, (**G**) 2.5 μg ml^-1^ daptomycin, (**H**) 5 μg ml^-1^ vancomycin, (**I**) 2.5 μg ml^-1^ gentamicin, (**J**) 2.5 μg ml^-1^ tetracycline, (**K**) 0.038 μg ml^-1^ rifampicin and (**L**) 40 μg ml^-1^ chloramphenicol. Data represent the geometric mean ± geometric standard deviation of three independent experiments. Data were analysed by one-way ANOVA with Dunnett’s *post-hoc* test (* P < 0.05, WT vs mutant).

### Lack of XerC sensitises *S. aureus* to immune-mediated killing

In the host, *S. aureus* DNA damage is not only caused by antibiotic exposure, but also by the immune system, particularly by reactive oxygen species produced by neutrophils. Therefore, we used a whole human blood model to test whether XerC was required for surviving the DNA damage since previous work from our group and others have shown that *S. aureus* in blood is rapidly phagocytosed by neutrophils and exposed to the oxidative burst [Ha *et al*., 2020; Painter *et al*., 2017].

The WT strain demonstrated high levels of survival on exposure to whole blood from healthy human donors, with 80 % of cells viable after 4 h (Fig. 6). By contrast, significantly lower survival (24 %) of the *xerC*::Tn mutant strain was observed (Fig. 6).

**Figure 5.**
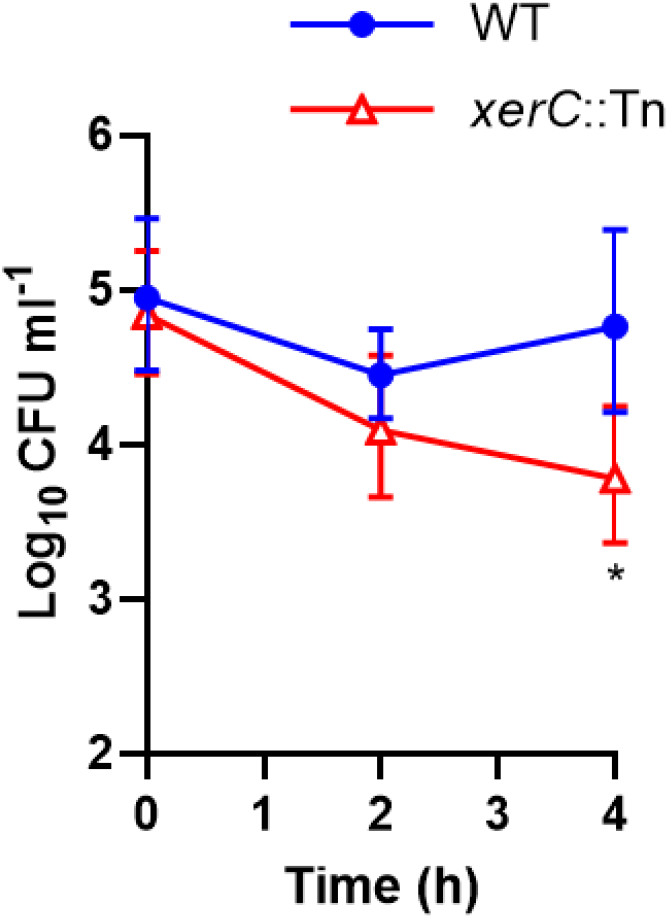
Lack of XerC sensitises *S. aureus* to killing by human blood. Log_10_ CFU ml^-1^ of *S. aureus* USA300 JE2 WT and the *xerC*::Tn mutant during a 4 h incubation in whole human blood. Data represent the geometric mean ± geometric standard deviation of three independent experiments. Data were analysed by one-way ANOVA with Dunnett’s *post-hoc* test (* P < 0.05, WT vs mutant).

Taken together, XerC is required for induction of the SOS response, with the result that a mutant defective in *xerC* is more susceptible to a range of DNA-damaging antibiotics and immune-mediated killing.

## Discussion

Treatment of infections caused by *S. aureus* is challenging due to increasing rates of antibiotic resistance. In addition to discovering novel antibiotics, an alternative approach is to develop compounds which sensitise bacteria to existing antibiotics. One such approach is to sensitise *S. aureus* to DNA-damaging antibiotics by inhibiting bacterial DNA repair via targeting the SOS response pathway [Lanyon-Hogg *et al*., 2021]. Unfortunately, this approach has been hindered by *S. aureus* having a poorly characterised SOS response, meaning that the best proteins to target have not yet been identified. Therefore, to address this, we measured the ability of 54 mutants in various DNA repair pathways to induce the SOS response on exposure to ciprofloxacin.

In *S. aureus*, DNA double strand breaks are processed by the RexAB enzyme complex to generate a piece of single-stranded DNA that is bound by RecA, resulting in the autocleavage of the LexA repressor and activation of the SOS response [Ha *et al*., 2020]. In agreement with this, our screen of mutants defective for various DNA repair proteins identified that lack of RexB significantly reduced SOS induction, validating our screening approach. Through our screen, we identified two additional genes, *xerC* and *xseA*, where disruption reduced SOS induction and increased ciprofloxacin susceptibility. *xseA*, together with *xseB* (not included in the screen), encodes DNA exonuclease VII which degrades single stranded DNA [Lovett 2011]. In *E. coli*, it has been suggested that this complex processes DNA double strand breaks to generate blunt DNA ends which can then be recognised by RecBCD, triggering the SOS response [Repar *et al*., 2013]. *xerC* encodes a tyrosine recombinase which, together with *xerD* (not included in the screen), is important for resolving the DNA multimers which form during DNA replication and repair [Blakely *et al*., 1991]. The reason why XerC is required for optimal induction of the SOS response in *S. aureus* is currently unclear, and contrasts with work in *E. coli* demonstrating that the absence of *xerC* did not affect SOS induction [Hendricks *et al*., 2000]. However, this finding further emphasises differences in the mechanisms of induction of the SOS response between *S. aureus* and much more well-characterised organisms such as *E. coli*.

As well as *xseA, xerC* and *rexB*, we also identified 15 genes where disruption significantly reduced SOS induction, but had no effect on ciprofloxacin susceptibility. In these cases, the SOS response was not as impaired as the mutants where DNA repair was inhibited, and so although the SOS response was reduced, there may have been a sufficient level to enable DNA repair. Alternatively, these mutants may upregulate alternative DNA repair pathways to compensate for the lack of SOS response induction. This finding has implications for the design of screens to identify new inhibitors of DNA repair, demonstrating the importance of characterising susceptibility to antibiotics as well as the SOS response.

In addition to resolving chromosomal multimers, XerC plays a key role in monomerising plasmids in many bacterial species including *E. coli, Salmonella typhimurium, Klebsiella pneumoniae*, and *Acinetobacter baumannii* [Lin *et al*., 2020; Sherratt *et al*., 1995; Bui *et al*., 2006]. Our finding that XerC was required for preventing plasmid loss in *S. aureus* indicates that XerC may play a similar role in this bacterium. This may have important implications for the spread of antibiotic resistance and virulence genes, as in *S. aureus* these are frequently located on plasmids [Haaber *et al*., 2017]. However, a limitation of this work is that these plasmid segregation assays were performed with an *E. coli*/*S. aureus* shuttle vector which may not be representative of plasmids found in clinical *S. aureus* strains. However, while large plasmids typically encode their own machinery for monomerisation, smaller plasmids found in clinical *S. aureus* strains do not, and instead rely on bacterially-encoded systems [Castillo *et al*., 2017]. Therefore, in some cases, therapeutic inhibition of XerC may lead to a loss of antibiotic resistance, sensitising *S. aureus* to antibiotics.

Beyond the loss of plasmid-mediated resistance, very few studies have examined the direct role of XerC in antibiotic susceptibility, especially in *S. aureus*. A previous screen of the NARSA library identified the mutant defective for *xerC* was more susceptible to ciprofloxacin and gentamicin but not to oxacillin, linezolid, fosfomycin, daptomycin, mupirocin or vancomycin [Vestergaard *et al*., 2016]. Increased ciprofloxacin susceptibility has also been reported for *E. coli* and *P. aeruginosa xerC* mutants [D<u>ö</u>rr *et al*., 2009; Baggett *et al*., 2021]. However, our finding that the *S. aureus xerC* mutant is more susceptible to antibiotics from a range of classes fits with the growing evidence that many antibiotics lead to DNA damage via the production of ROS [Dwyer *et al*., 2014]. The reason why XerC is required for repair of antibiotic induced DNA damage is currently unknown. However, one hypothesis is that when bacteria repair DNA damage through homologous recombination, DNA dimers are produced which must be resolved by the activity of XerCD to enable subsequent DNA replication to occur and for correct DNA segregation during cell division [Castillo *et al*., 2017]. However, this remains to be demonstrated experimentally.

Finally, in agreement with data showing that neutrophils cause DNA double strand breaks which are repaired via the SOS response, the *xerC*::Tn mutant showed significantly reduced survival in a whole human blood model [Ha *et al*., 2020]. This adds to our appreciation of the importance of DNA repair pathways inside the host and provides a possible explanation for the reduced virulence of the *xerC* mutants observed in murine bacteraemia and pyelonephritis models [Atwood *et al*., 2016; Chalker *et al*., 2000]. Additionally, it may suggest that chemical inhibition of XerC may be a viable monotherapeutic approach as it may enhance bacterial killing by the immune system.

In summary, characterisation of the SOS response in *S. aureus* has led to the identification of XerC as a key protein required for the induction of this pathway. The benefits of targeting this protein complex are numerous as it is required for survival inside the host, meaning that inhibitors of XerC may have activity alone, reducing bacterial virulence and enhancing immune-mediated killing. In addition, they may potentiate various classes of existing antibiotic, including directly DNA damaging antibiotics such as ciprofloxacin and co-trimoxazole, but also cell wall targeting agents and protein synthesis inhibitors. Finally, inhibitors of XerC may reduce rates of antibiotic resistance by preventing the segregation of plasmids carrying antibiotic resistance genes, or by blocking the SOS response. As well as reducing the mutation rate and therefore the emergence of spontaneous resistance, this also suppresses the activation of prophages, slowing the dissemination of antibiotic resistance and virulence genes.

## Materials and methods

### Bacterial strains and growth conditions

The bacterial strains used in this study are shown in Table S2. *S. aureus* was grown in tryptic soy broth (TSB) for 16 h at 37 °C with shaking (180 rpm). When required, TSB was supplemented with erythromycin (10 μg ml^-1^), kanamycin (90 μg ml^-1^) or chloramphenicol (5 μg ml^-1^). For strains complemented with p*itet*, media were supplemented with 100 ng ml^-1^ anhydrotetracycline (Atc) to induce expression.

### Construction of strains

The P*recA-gfp* reporter plasmid was transduced from JE2 WT P*recA-gfp* [Clarke *et al*., 2019] into mutants obtained from the NARSA library using ϕ11, as previously described [Olsen, 2016].

Complementation of the *xerC*::Tn mutant was carried out using p*itet* [Gründling and Schneewind, 2007]. The *xerC* gene was amplified from JE2 WT genomic DNA using *xerC*_Fw and *xerC*_Rev primers (Table S3). To construct p*itet*-*xerC*, the PCR product and the p*itet* vector were digested with avrII and pmeI, ligated using T4 ligase and transformed into *E. coli* DC10B. p*itet-xerC* was then electroporated into RN4220 and then transduced into the *xerC*::Tn mutant using ϕ11. To construct *xerC*::Tn p*itet-xerC*^Y273F^, site directed mutagenesis was performed using p*itet-xerC* as a template, the *xerC-*Y273F_Fw and *xerC*-Y273F_Rev primers (Table S3) and the Q5 site directed mutagenesis kit according to the manufacturer’s instructions. From *E. coli*, p*itet-xerC*^Y273F^ was transformed into RN4220 and then transduced into the *xerC*::Tn mutant using ϕ11.

### Measurement of SOS response by fluorescent reporter assay

Strains containing the P*recA-gfp* reporter plasmid were used to quantify *recA* expression as a measure of induction of the SOS response as previously described [Clarke *et al*., 2019]. Two-fold serial dilutions of antibiotic were performed in TSB in black 96-well plates in a final volume of 200 μl. Wells were inoculated with 10^8^ CFU ml^-1^ and plates were incubated for 17 h in a Tecan Infinite M200-PRO plate reader at 37°C with orbital shaking (300 rpm). Every 15 min, fluorescence (excitation 485 nm; emission 525 nm) and OD_600_ were measured. RFU values were divided by OD_600_ at each time point to normalise for changes in cell density which occurred during the assay and the peak RFU/OD_600_ was plotted.

### Measurements of plasmid stability

To determine the rate at which the P*recA-gfp* reporter plasmid was lost from the JE2 WT and *xerC*::Tn mutant strains each of these strains containing the plasmid were passaged six times in the presence or absence of kanamycin selection. Each passage consisted of a 1000-fold dilution in TSB supplemented, or not, with 90 μg ml^-1^ kanamycin and then 8 h growth at 37°C with shaking (180 rpm). Cultures were stored overnight between passages at 4°C. After each passage, each culture was serially diluted 10-fold in PBS and plated onto TSA with and without 90 μg ml^-1^ kanamycin to enumerate CFU ml^-1^. The CFU ml^-1^ on TSA + kanamycin was divided by the CFU ml^-1^ on TSA to calculate the proportion of CFU containing the plasmid.

### Determination of antibiotic minimum inhibitory concentrations

MICs were determined using a broth microdilution protocol as described previously [Wiegand *et al*., 2008]. Two-fold serial dilutions of antibiotics were prepared in a final volume of 200 μl TSB in 96 well plates. For daptomycin, 1.25 mM CaCl_2_ was added to the media. Wells were inoculated to 5 × 10^5^ CFU ml^-1^ bacteria and incubated statically for 17 h at 37 °C. The MIC was defined as the minimum concentration of antibiotic where no visible growth was observed.

### Antibiotic killing assay

TSB (3 ml) supplemented with antibiotics at 5 x WT MIC was inoculated with 10^8^ CFU ml^-1^ *S. aureus*. Cultures were incubated at 37 for 6 h with shaking (180 rpm). For daptomycin, 1.25 mM CaCl_2_ was added to the media. After 0 h and 6 h antibiotic exposure, cultures were serially diluted 10-fold in PBS and plated on TSA to enumerate CFU ml^-1^.

### Whole human blood killing assay

Ethical approval for drawing and using human blood was obtained from the Regional Ethics Committee and the Imperial NHS Trust Tissue Bank (REC Wales approval no. 12/WA/0196 and ICHTB HTA license no. 12275). Whole human blood was collected in EDTA tubes. Overnight cultures of *S. aureus* were washed twice in PBS and resuspended at 10^6^ CFU ml^-1^. In a 96 well plate, 90 μl blood was inoculated with 10 μl bacteria to give a final inoculum of 10^5^ CFU ml^-1^. Plates were incubated at 37°C with shaking (180 rpm) and after 0, 2 and 4 h, samples were serially diluted 10-fold in PBS and plated onto TSA to enumerate surviving CFU ml^-1^.

### Statistical analyses

CFU data were log_10_ transformed and presented as the geometric mean ± geometric standard deviation. Non-CFU data were presented as the mean ± standard deviation, except MIC data which were presented as the median value. All experiments consisted of at least three independent biological replicates and were analysed as described in the figure legends using GraphPad Prism (V8.0).

## Acknowledgements

EVKL, EWT and AME acknowledge support from the Rosetrees Trust. AME acknowledges support from the Imperial NIHR Biomedical Research Centre, Imperial College London.

All authors acknowledge the provision of strains by the Network on Antimicrobial Resistance in *Staphylococcus aureus* (NARSA) Program: under NIAID/ NIH Contract No. HHSN272200700055C. The funders had no role in the study design, interpretation of the findings or the writing of the manuscript.

