## Supplementary data for "XerC is required for the repair of antibiotic- and immune-mediated DNA damage in *Staphylococcus aureus*"

**Supplementary information**

**Supplementary Figures 1-2**

**Supplementary Tables 1-3**

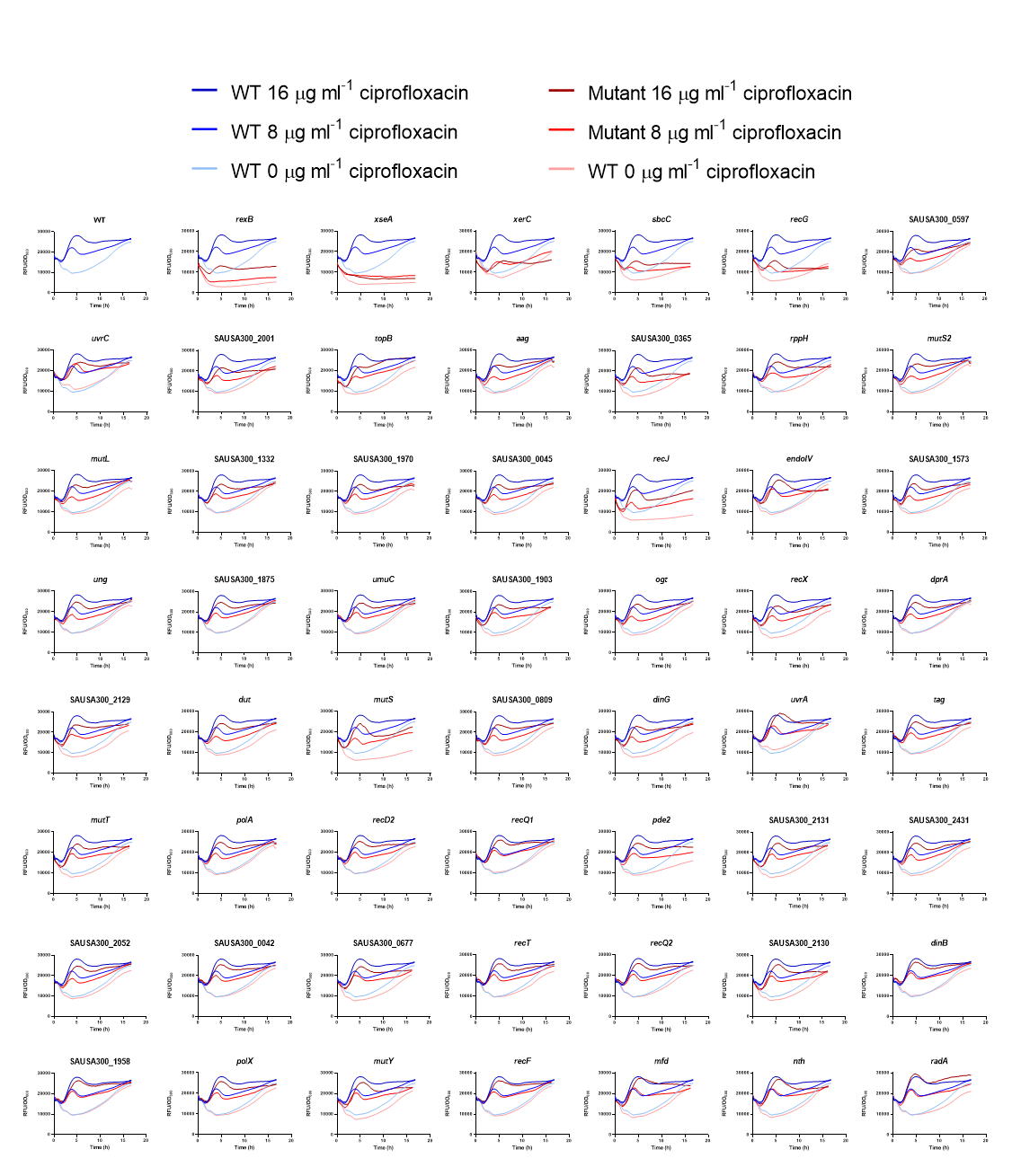
**Fig. S1. SOS response of DNA repair mutants.** USA300 *S. aureus* WT and NARSA mutants defective for various DNA repair genes containing the P*recA*-*gfp* reporter were exposed to 0, 8 or 16 µg ml^-1^ ciprofloxacin and GFP and OD_600_ measured over 17 h. GFP was divided by OD_600_ to normalise for changes in cell density which occurred during the assay.

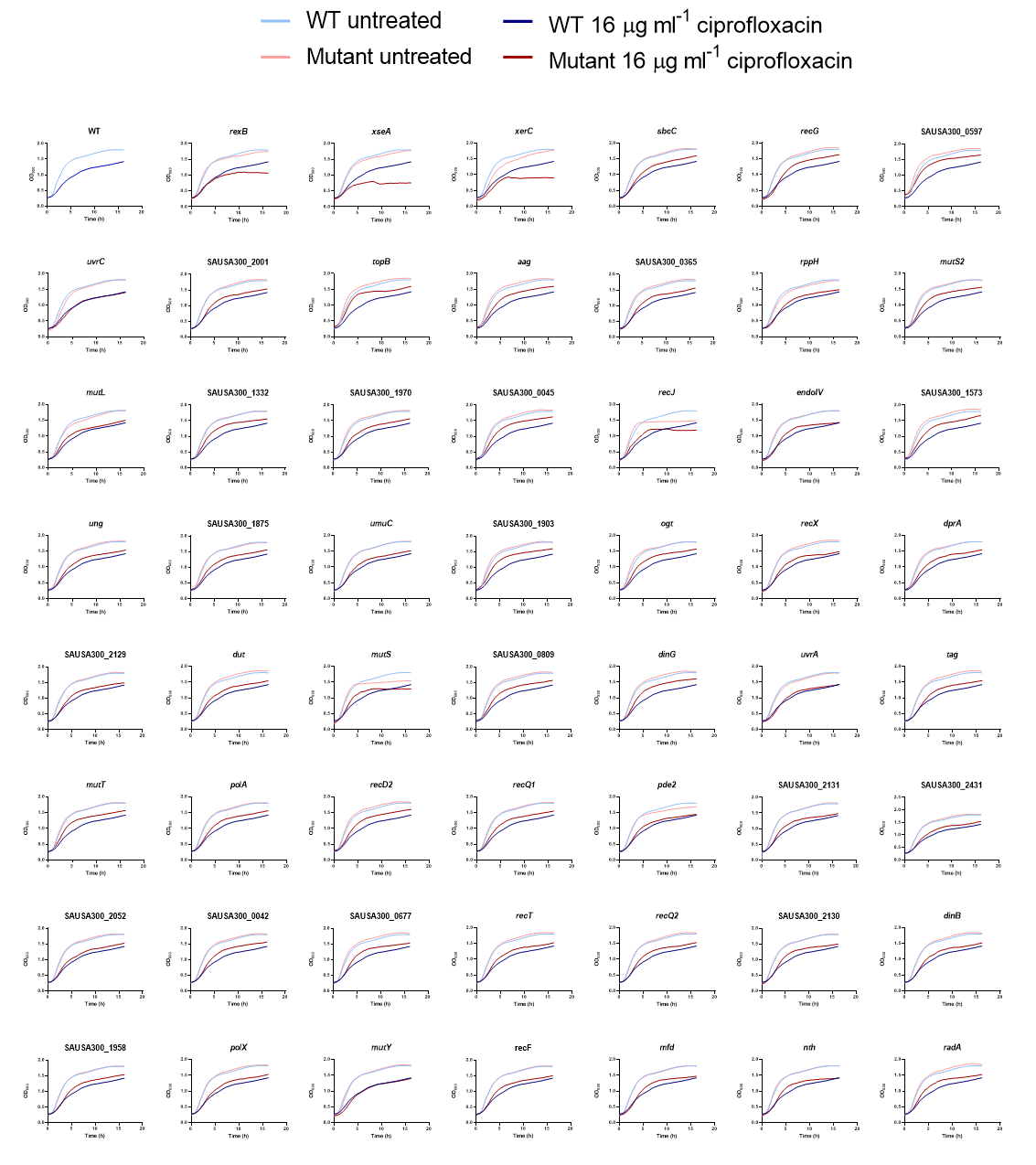
**Fig. S2. Ciprofloxacin susceptibility of DNA repair mutants.** USA300 *S. aureus* WT and NARSA mutants defective for various DNA repair genes containing the P*recA*-*gfp* reporter were exposed to 0 or 16 µg ml^-1^ ciprofloxacin and OD_600_ measured over 17 h.

**Table S1. Mutants screened for ability to induce SOS response**

| NARSA reference | Gene | Gene name | Description |
| --- | --- | --- | --- |
| NE1012 | SAUSA300_0869 | *rexB* | Exonuclease RexB |
| NE458 | SAUSA300_1472 | *xseA* | Exodeoxyribonuclease VII, large subunit |
| NE1451 | SAUSA300_1243 | *sbcC* | Exonuclease |
| NE1344 | SAUSA300_1120 | *recG* | ATP-dependent DNA helicase |
| NE883 | SAUSA300_1145 | *xerC* | Tyrosine recombinase XerC |
| NE1878 | SAUSA300_0597 |  | Putative endonuclease III |
| NE1212 | SAUSA300_1045 | *uvrC* | Excinuclease ABC subunit C |
| NE93 | SAUSA300_2001 |  | Similar to DNA mismatch repair protein |
| NE152 | SAUSA300_2208 | *topB* | DNA topoisomerase III |
| NE1613 | SAUSA300_2290 | *aag* | Putative 3-methyladenine DNA glycosylase |
| NE1900 | SAUSA300_0365 |  | Hypothetical protein |
| NE746 | SAUSA300_1734 | *rppH* | Conserved hypothetical protein |
| NE1462 | SAUSA300_1043 | *mutS2* | DNA mismatch repair MutS2 protein |
| NE80 | SAUSA300_1189 | *mutL* | DNA mismatch repair protein MutL |
| NE789 | SAUSA300_1332 |  | Putative 5'-3' exonuclease |
| NE246 | SAUSA300_1970 |  | Putative exonuclease |
| NE88 | SAUSA300_0045 |  | HNH endonuclease family protein |
| NE11 | SAUSA300_1592 | *recJ* | ssDNA exonuclease |
| NE1028 | SAUSA300_1517 | *endoIV* | Endonuclease IV |
| NE1794 | SAUSA300_1573 |  | Holliday junction resolvase-like protein |
| NE888 | SAUSA300_0563 | *ung* | Uracil-DNA glycosylase |
| NE1320 | SAUSA300_1875 |  | Exonuclease |
| NE445 | SAUSA300_1259 | *umuC* | ImpB/MucB/SamB family protein |
| NE692 | SAUSA300_1903 |  | Conserved hypothetical protein |
| NE1907 | SAUSA300_2485 | *ogt* | Methylated DNA-protein cysteine methyltransferase |
| NE324 | SAUSA300_1854 | *recX* | Regulatory protein RecX |
| NE242 | SAUSA300_1142 | *dprA* | DNA protecting protein DprA |
| NE993 | SAUSA300_2129 |  | Putative hemolysin III (regulated by RecA) |
| NE1798 | SAUSA300_1949 | *dut* | dUTP diphosphatase |
| NE974 | SAUSA300_1188 | *mutS* | DNA mismatch repair protein mutS |
| NE466 | SAUSA300_0809 |  | Putative DNA primase |
| NE346 | SAUSA300_1346 | *dinG* | Putative DnaQ family exonuclease/DinG family helicase |
| NE145 | SAUSA300_0742 | *uvrA* | Excinuclease ABC, A subunit |
| NE1825 | SAUSA300_1612 | *tag* | DNA-3-methyladenine glycosidase |
| NE1653 | SAUSA300_2432 | *mutT* | MutT/NUDIX family hydrolase |
| NE22 | SAUSA300_1636 | *polA* | DNA polymerase I superfamily |
| NE1427 | SAUSA300_1576 | *recD2* | Helicase, RecD/TraA family |
| NE972 | SAUSA300_0705 | *recQ1* | ATP-dependent DNA helicase RecQ |
| NE1208 | SAUSA300_1650 | *pde2* | Hypothetical protein |
| NE1146 | SAUSA300_2131 |  | Hypothetical protein (regulated by RecA) |
| NE513 | SAUSA300_2431 |  | Putative helicase |
| NE1679 | SAUSA300_2052 |  | Single-stranded DNA- binding protein family |
| NE487 | SAUSA300_0042 |  | Conserved hypothetical protein |
| NE279 | SAUSA300_0677 | *phrB* | Putative DNA photolyase (regulated by RecA) |
| NE1830 | SAUSA300_1960 | *recT* | Putative phage-related DNA recombination protein |
| NE1528 | SAUSA300_1371 | *recQ2* | ATP-dependent DNA helicase RecQ |
| NE97 | SAUSA300_2130 |  | UTP-glucose-1-phosphate uridylyltransferase family protein (regulated by RecA) |
| NE1866 | SAUSA300_1876 | *dinB* | DNA polymerase IV |
| NE824 | SAUSA300_1958 |  | Single-strand binding protein |
| NE947 | SAUSA300_1042 | *polX* | Hypothetical protein |
| NE1040 | SAUSA300_1849 | *mutY* | A/G-specific adenine glycosylase |
| NE555 | SAUSA300_0004 | *recF* | DNA replication and repair protein RecF |
| NE188 | SAUSA300_0481 | *mfd* | Transcription-repair coupling factor |
| NE761 | SAUSA300_1343 | *nth* | Endonuclease III |
| NE1176 | SAUSA300_0511 | *radA* | DNA repair protein RadA |

**Table S2. Strains used in this study**

| **Strain** | **Relevant characteristics** | **Reference/source** |
| --- | --- | --- |
| USA300 JE2 WT | LAC strain of the USA300 CA-MRSA lineage cured of plasmids | Fey *et al.,* 2013 |
| USA300 JE2 *xerC*::Tn | JE2 with a *bursa aurealis* transposon insertion in *xerC*, Ery^r^ | Fey *et al.,* 2013 |
| USA300 JE2 *xerC*::Tn p*itet-xerC* | JE2 with a *bursa aurealis* transposon insertion in *xerC* complemented with *pitet-xerC.* Ery*^r^,* Cm^r^ | This study |
| USA300 JE2 *xerC*::Tn p*itet-xerC*^Y273F^ | JE2 with a *bursa aurealis* transposon insertion in *xerC* complemented with *pitet-xerC*^Y273F^*.* Ery*^r^,* Cm^r^. | This study |
| USA300 JE2 WT P*recA*_*gfp* | USA300 LAC JE2 carrying the P*recA*-*gfp* reporter plasmid, Kan^r^ | Ha *et al.,* 2020 |
| USA300 JE2 *rexB*::Tn P*recA_gfp* | JE2 with a *bursa aurealis* transposon insertion in *rexB* carrying the P*recA*-*gfp* reporter plasmid, Kan^r^, Ery*^r^.* | Ha *et al.,* 2020 |
| USA300 JE2 *uvrA*::Tn P*recA_gfp* | JE2 with a *bursa aurealis* transposon insertion in *uvrA* carrying the P*recA*-*gfp* reporter plasmid, Kan^r^, Ery*^r^* | This study |
| USA300 JE2 *sbcC*::Tn P*recA_gfp* | JE2 with a *bursa aurealis* transposon insertion in *sbcC* carrying the P*recA*-*gfp* reporter plasmid, Kan^r^, Ery*^r^* | This study |
| USA300 JE2 *recF*::Tn P*recA_gfp* | JE2 with a *bursa aurealis* transposon insertion in *recF* carrying the P*recA*-*gfp* reporter plasmid, Kan^r^, Ery*^r^* | This study |
| USA300 JE2 *nth*::Tn P*recA_gfp* | JE2 with a *bursa aurealis* transposon insertion in *nth* carrying the P*recA*-*gfp* reporter plasmid, Kan^r^, Ery*^r^* | This study |
| USA300 JE2 *uvrC*::Tn P*recA_gfp* | JE2 with a *bursa aurealis* transposon insertion in *uvrC* carrying the P*recA*-*gfp* reporter plasmid, Kan^r^, Ery*^r^* | This study |
| USA300 JE2 *mutS*::Tn P*recA_gfp* | JE2 with a *bursa aurealis* transposon insertion in *mutS* carrying the P*recA*-*gfp* reporter plasmid, Kan^r^, Ery*^r^* | This study |
| USA300 JE2 *recJ*::Tn P*recA_gfp* | JE2 with a *bursa aurealis* transposon insertion in *recJ* carrying the P*recA*-*gfp* reporter plasmid, Kan^r^, Ery*^r^* | This study |
| USA300 JE2 *endoIV*::Tn P*recA_gfp* | JE2 with a *bursa aurealis* transposon insertion in *endoIV* carrying the P*recA*-*gfp* reporter plasmid, Kan^r^, Ery*^r^* | This study |
| USA300 JE2 *xerC*::Tn P*recA_gfp* | JE2 with a *bursa aurealis* transposon insertion in *xerC* carrying the P*recA*-*gfp* reporter plasmid, Kan^r^, Ery*^r^* | This study |
| USA300 JE2 *recG*::Tn P*recA_gfp* | JE2 with a *bursa aurealis* transposon insertion in *recG* carrying the P*recA*-*gfp* reporter plasmid, Kan^r^, Ery*^r^* | This study |
| USA300 JE2 *mutT*::Tn P*recA_gfp* | JE2 with a *bursa aurealis* transposon insertion in *mutT* carrying the P*recA*-*gfp* reporter plasmid, Kan^r^, Ery*^r^* | This study |
| USA300 JE2 *mutY*::Tn P*recA_gfp* | JE2 with a *bursa aurealis* transposon insertion in *mutY* carrying the P*recA*-*gfp* reporter plasmid, Kan^r^, Ery*^r^* | This study |
| USA300 JE2 SAUSA300_2130::Tn P*recA_gfp* | JE2 with a *bursa aurealis* transposon insertion in SAUSA300_2130 carrying the P*recA*-*gfp* reporter plasmid, Kan^r^, Ery*^r^* | This study |
| USA300 JE2 SAUSA300_0677::Tn P*recA_gfp* | JE2 with a *bursa aurealis* transposon insertion in SAUSA300_0677 carrying the P*recA*-*gfp* reporter plasmid, Kan^r^, Ery*^r^* | This study |
| USA300 JE2 SAUSA300_1903::Tn P*recA_gfp* | JE2 with a *bursa aurealis* transposon insertion in SAUSA300_1903 carrying the P*recA*-*gfp* reporter plasmid, Kan^r^, Ery*^r^* | This study |
| USA300 JE2 SAUSA300_2131::Tn P*recA_gfp* | JE2 with a *bursa aurealis* transposon insertion in SAUSA300_2131 carrying the P*recA*-*gfp* reporter plasmid, Kan^r^, Ery*^r^* | This study |
| USA300 JE2 SAUSA300_0365::Tn P*recA_gfp* | JE2 with a *bursa aurealis* transposon insertion in SAUSA300_0365 carrying the P*recA*-*gfp* reporter plasmid, Kan^r^, Ery*^r^* | This study |
| USA300 JE2 *mfd*::Tn P*recA_gfp* | JE2 with a *bursa aurealis* transposon insertion in *mfd* carrying the P*recA*-*gfp* reporter plasmid, Kan^r^, Ery*^r^* | This study |
| USA300 JE2 SAUSA300_2129::Tn P*recA_gfp* | JE2 with a *bursa aurealis* transposon insertion in SAUSA300_2129 carrying the P*recA*-*gfp* reporter plasmid, Kan^r^, Ery*^r^* | This study |
| USA300 JE2 *ogt*::Tn P*recA_gfp* | JE2 with a *bursa aurealis* transposon insertion in *ogt* carrying the P*recA*-*gfp* reporter plasmid, Kan^r^, Ery*^r^* | This study |
| USA300 JE2 *ung*::Tn P*recA_gfp* | JE2 with a *bursa aurealis* transposon insertion in *ung* carrying the P*recA*-*gfp* reporter plasmid, Kan^r^, Ery*^r^* | This study |
| USA300 JE2 *aag*::Tn P*recA_gfp* | JE2 with a *bursa aurealis* transposon insertion in *aag* carrying the P*recA*-*gfp* reporter plasmid, Kan^r^, Ery*^r^* | This study |
| USA300 JE2 SAUSA300_0809::Tn P*recA_gfp* | JE2 with a *bursa aurealis* transposon insertion in SAUSA300_0809 carrying the P*recA*-*gfp* reporter plasmid, Kan^r^, Ery*^r^* | This study |
| USA300 JE2 *polA*::Tn P*recA_gfp* | JE2 with a *bursa aurealis* transposon insertion in *polA* carrying the P*recA*-*gfp* reporter plasmid, Kan^r^, Ery*^r^* | This study |
| USA300 JE2 *dut*::Tn P*recA_gfp* | JE2 with a *bursa aurealis* transposon insertion in *dut* carrying the P*recA*-*gfp* reporter plasmid, Kan^r^, Ery*^r^* | This study |
| USA300 JE2 *recJ*::Tn P*recA_gfp* | JE2 with a *bursa aurealis* transposon insertion in *recJ* carrying the P*recA*-*gfp* reporter plasmid, Kan^r^, Ery*^r^* | This study |
| USA300 JE2 *mutS2*::Tn P*recA_gfp* | JE2 with a *bursa aurealis* transposon insertion in *mutS2* carrying the P*recA*-*gfp* reporter plasmid, Kan^r^, Ery*^r^* | This study |
| USA300 JE2 *pde2*::Tn P*recA_gfp* | JE2 with a *bursa aurealis* transposon insertion in *pde2* carrying the P*recA*-*gfp* reporter plasmid, Kan^r^, Ery*^r^* | This study |
| USA300 JE2 *recQ1*::Tn P*recA_gfp* | JE2 with a *bursa aurealis* transposon insertion in *recQ1* carrying the P*recA*-*gfp* reporter plasmid, Kan^r^, Ery*^r^* | This study |
| USA300 JE2 *recT*::Tn P*recA_gfp* | JE2 with a *bursa aurealis* transposon insertion in *recT* carrying the P*recA*-*gfp* reporter plasmid, Kan^r^, Ery*^r^* | This study |
| USA300 JE2 *rppH*::Tn P*recA_gfp* | JE2 with a *bursa aurealis* transposon insertion in *rppH* carrying the P*recA*-*gfp* reporter plasmid, Kan^r^, Ery*^r^* | This study |
| USA300 JE2 *recD2*::Tn P*recA_gfp* | JE2 with a *bursa aurealis* transposon insertion in *recD2* carrying the P*recA*-*gfp* reporter plasmid, Kan^r^, Ery*^r^* | This study |
| USA300 JE2 *recQ2*::Tn P*recA_gfp* | JE2 with a *bursa aurealis* transposon insertion in *recQ2* carrying the P*recA*-*gfp* reporter plasmid, Kan^r^, Ery*^r^* | This study |
| USA300 JE2 *polX*::Tn P*recA_gfp* | JE2 with a *bursa aurealis* transposon insertion in *polX* carrying the P*recA*-*gfp* reporter plasmid, Kan^r^, Ery*^r^* | This study |
| USA300 JE2 *mutL*::Tn P*recA_gfp* | JE2 with a *bursa aurealis* transposon insertion in *mutL* carrying the P*recA*-*gfp* reporter plasmid, Kan^r^, Ery*^r^* | This study |
| USA300 JE2 *dinG*::Tn P*recA_gfp* | JE2 with a *bursa aurealis* transposon insertion in *dinG* carrying the P*recA*-*gfp* reporter plasmid, Kan^r^, Ery*^r^* | This study |
| USA300 JE2 SAUSA300_2431::Tn P*recA_gfp* | JE2 with a *bursa aurealis* transposon insertion in SAUSA300_2431 carrying the P*recA*-*gfp* reporter plasmid, Kan^r^, Ery*^r^* | This study |
| USA300 JE2 SAUSA300_1573::Tn P*recA_gfp* | JE2 with a *bursa aurealis* transposon insertion in SAUSA300_1573 carrying the P*recA*-*gfp* reporter plasmid, Kan^r^, Ery*^r^* | This study |
| USA300 JE2 SAUSA300_0597::Tn P*recA_gfp* | JE2 with a *bursa aurealis* transposon insertion in SAUSA300_0597 carrying the P*recA*-*gfp* reporter plasmid, Kan^r^, Ery*^r^* | This study |
| USA300 JE2 SAUSA300_2502::Tn P*recA_gfp* | JE2 with a *bursa aurealis* transposon insertion in SAUSA300_2502 carrying the P*recA*-*gfp* reporter plasmid, Kan^r^, Ery*^r^* | This study |
| USA300 JE2 SAUSA300_1958::Tn P*recA_gfp* | JE2 with a *bursa aurealis* transposon insertion in SAUSA300_1958 carrying the P*recA*-*gfp* reporter plasmid, Kan^r^, Ery*^r^* | This study |
| USA300 JE2 *dinB*::Tn P*recA_gfp* | JE2 with a *bursa aurealis* transposon insertion in *dinB* carrying the P*recA*-*gfp* reporter plasmid, Kan^r^, Ery*^r^* | This study |
| USA300 JE2 *umuC*::Tn P*recA_gfp* | JE2 with a *bursa aurealis* transposon insertion in *umuC* carrying the P*recA*-*gfp* reporter plasmid, Kan^r^, Ery*^r^* | This study |
| USA300 JE2 *recX*::Tn P*recA_gfp* | JE2 with a *bursa aurealis* transposon insertion in *recX* carrying the P*recA*-*gfp* reporter plasmid, Kan^r^, Ery*^r^* | This study |
| USA300 JE2 SAUSA300_1875::Tn P*recA_gfp* | JE2 with a *bursa aurealis* transposon insertion in SAUSA300_1875 carrying the P*recA*-*gfp* reporter plasmid, Kan^r^, Ery*^r^* | This study |
| USA300 JE2 *dprA*::Tn P*recA_gfp* | JE2 with a *bursa aurealis* transposon insertion in *dprA* carrying the P*recA*-*gfp* reporter plasmid, Kan^r^, Ery*^r^* | This study |
| USA300 JE2 SAUSA300_2001::Tn P*recA_gfp* | JE2 with a *bursa aurealis* transposon insertion in SAUSA300_2001 carrying the P*recA*-*gfp* reporter plasmid, Kan^r^, Ery*^r^* | This study |
| USA300 JE2 *topB*::Tn P*recA_gfp* | JE2 with a *bursa aurealis* transposon insertion in *topB* carrying the P*recA*-*gfp* reporter plasmid, Kan^r^, Ery*^r^* | This study |
| USA300 JE2 SAUSA300_1332::Tn P*recA_gfp* | JE2 with a *bursa aurealis* transposon insertion in SAUSA300_1332 carrying the P*recA*-*gfp* reporter plasmid, Kan^r^, Ery*^r^* | This study |
| USA300 JE2 SAUSA300_1955::Tn P*recA_gfp* | JE2 with a *bursa aurealis* transposon insertion in SAUSA300_1955 carrying the P*recA*-*gfp* reporter plasmid, Kan^r^, Ery*^r^* | This study |
| USA300 JE2 SAUSA300_0042::Tn P*recA_gfp* | JE2 with a *bursa aurealis* transposon insertion in SAUSA300_0042 carrying the P*recA*-*gfp* reporter plasmid, Kan^r^, Ery*^r^* | This study |
| USA300 JE2 *radA*::Tn P*recA_gfp* | JE2 with a *bursa aurealis* transposon insertion in *radA* carrying the P*recA*-*gfp* reporter plasmid, Kan^r^, Ery*^r^* | This study |
| USA300 JE2 SAUSA300_1970::Tn P*recA_gfp* | JE2 with a *bursa aurealis* transposon insertion in SAUSA300_1970 carrying the P*recA*-*gfp* reporter plasmid, Kan^r^, Ery*^r^* | This study |
| USA300 JE2 SAUSA300_0045::Tn P*recA_gfp* | JE2 with a *bursa aurealis* transposon insertion in SAUSA300_0045 carrying the P*recA*-*gfp* reporter plasmid, Kan^r^, Ery*^r^* | This study |
| USA300 JE2 *tag*::Tn P*recA_gfp* | JE2 with a *bursa aurealis* transposon insertion in *tag* carrying the P*recA*-*gfp* reporter plasmid, Kan^r^, Ery*^r^* | This study |
| USA300 JE2 *xseA*::Tn P*recA_gfp* | JE2 with a *bursa aurealis* transposon insertion in *xseA* carrying the P*recA*-*gfp* reporter plasmid, Kan^r^, Ery*^r^* | This study |

**Table S3. Primers used in this study.**

| **Primer** | **Sequence (5’ – 3’) –** restriction sites underlined |
| --- | --- |
| *xerC*_Fw | ATGCCCTAGGGTATTGAATCATATTCAAGATGCG |
| *xerC*_Rev | CGATGTTTAAACGTATTACTCATGTTTCATTCTCC |
| *xerC*_Y273F_Fw | ACTGGTAAATTTACACACGTATC |
| *xerC_*Y273F_Rev | TGTTGACAAATTAACATGAC |
